# Quantitative comparison of CRISPR-Cas9-mediated mutation efficiency between mice and MEFs using digital PCR assays

**DOI:** 10.1101/2022.09.09.507282

**Authors:** Kwangjun Lee, Choogon Lee

**Affiliations:** Department of Biomedical Sciences, Program in Neuroscience, College of Medicine, Florida State University, 1115 West Call Street, Tallahassee, FL 32306, USA

**Keywords:** CRISPR, circadian, digital PCR, mutant mouse, MEFs, Period gene

## Abstract

The creation of mutant mice has been invaluable for advancing biomedical science, but is too time- and resource-intensive for investigating the full range of mutations and polymorphisms. Cell culture models are therefore an invaluable complement to mouse models, especially for cell-autonomous pathways like the circadian clock. In this study, we quantitatively assessed the use of CRISPR to create cell models in MEFs as compared to mouse models. We generated two point mutations in the clock genes *Per1* and *Per2* in mice and in MEFs using the same sgRNAs and repair templates for HDR and quantified the frequency of the mutations by digital PCR. The frequency was about an order of magnitude higher in mouse zygotes compared to that in MEFs. However, the mutation frequency in MEFs was still high enough for clonal isolation by simple screening of a few dozen individual cells. The *Per* mutant cells that we generated provide important new insights into the role of the PAS domain in regulating PER phosphorylation, a key aspect of the circadian clock mechanism. Accurate quantification of the mutation frequency in bulk MEF populations provides a critical basis for optimizing CRISPR protocols and time/resource planning for generating cell models for further studies.

## Introduction

The prokaryotic defense system CRISPR-Cas is an adaptive immunity against foreign DNA molecules and has been converted to a revolutionary genome editing tool^1-3^. The CRISPR-Cas9 system from *Streptococcus pyogenes* is the most widely used because of its efficiency and simplicity^4^. The system requires two components: the Cas9 nuclease and a single guide RNA (sgRNA) which targets the nuclease to a specific genomic locus based on base pairing at a 20-nt target sequence. The sgRNA-guided Cas9 generates a double strand break (DSB) in the target site, which may be repaired in one of two different ways. The first is random insertion/deletion (indel) mutagenesis which occurs when the DSB is repaired by the error-prone non-homologous end-joining (NHEJ) pathway, resulting in a knockout allele due to frameshifting or indels of amino acids (AAs) due to in-frame nucleotide indels^4^. The second is precise allele editing through the high-fidelity homology-directed repair (HDR) pathway based on a donor or repair template, which usually occurs less frequently than NHEJ^5, 6^. More versatile than other genetic approaches such as siRNA and transgene expression, and more efficient than older homologous recombination methods^7, 8^, CRISPR is becoming a more and more common method to modulate a gene to interrogate its function or for medical applications^9, 10^.

CRISPR technology is rapidly evolving but still has many limitations. If target cells do not proliferate or are not easily transfected or electroporated with plasmids, clonal selection and expansion from the treated heterogenic population may not be possible or practical. CRISPR viral vectors have been employed to try to overcome these problems^11-16^. Clonal selection and expansion may not be required if the transduction efficiency and expression of the viral vectors are close to 100% and sgRNA efficiency is very high^17^. Several strategies have been developed to enhance HDR efficiency by activating HDR-related proteins or inhibiting NHEJ-related proteins ^18-20^. HDR donor templates also affect the efficiency significantly. Various lengths, chemical modifications, donor strand preference and symmetry of homologous arms have been tested ^4, 21-23^. Although results vary significantly in these studies, it seems that asymmetric single stranded oligodeoxynucleotides (ssODNs) with phosphorothioate bonds from the non-targeting strand are most effective as HDR donor templates.

The circadian clock drives daily rhythms in behavior and physiology ^24-28^, and dysfunction or disruption of the clock has been implicated in diverse disease states including sleep disorders ^29-33^. Decades of prior work have revealed that the clock is built on a core feedback loop that is cell autonomous, involving transcriptional and post-translational regulation of the redundant pacemaker *Period* (*Per*) genes *Per1* and *Per2*^34, 35^. Because the circadian clock is cell autonomous, genetic disruptions of the clock manifest similar phenotypes at the behavioral and cellular levels, and cell culture has proven to be a valuable and valid platform for characterizing the molecular biology of circadian rhythms ^36-38^. The endogenous clocks of cultured cells—including mouse embryonic fibroblasts (MEFs) and human U2OS cells—can be precisely measured in real time by introducing a luciferase (Luc) reporter gene under control of a clock promoter ^36, 38-40^. Across numerous studies, such cells have served as functional models for in vivo circadian clocks, and results have been consistently validated in live animal models. Cell culture models are not only less resource-consuming, but also more easily manipulated by chemicals and transgenes, which makes the cell models more suitable for mechanistic studies.

Historically, manipulation of endogenous clock genes in cell culture models suffered from technical limitations. Many genetic cell models required first developing mutant mouse models from which cells were then harvested. For example, mice with the *mPer2-Luc* knockin gene were crossed with mice with other mutations, backcrossed as needed, and MEFs were obtained from the resulting transgenic/mutant offspring ^41-43^. Recent developments in genome editing have created new opportunities for generating cell culture models without first generating mutant mice. Several studies including ours demonstrated that clock genes can be knocked out efficiently in culture using CRISPR^17, 44-47^. To our knowledge, however, there are no HDR-mediated mutations made in clock genes in MEFs by CRISPR. Because many mutant and transgenic mouse models including *mPer2-Luc* knockin are already available, it would be advantageous to implement direct genome editing in MEFs derived from these existing genetic mouse models.

In this current study we generated two CRISPR-mediated SNP mutations in mice and MEFs and quantified and compared the efficiency between mice and MEFs. Digital PCR can be a powerful tool for genotyping of CRISPR mutant mice when indels are too large to be detected by the conventional PCR-based genotyping. When HDR-mediated mutations are generated in cells, we show that digital PCR is a simple yet powerful tool to accurately quantify the frequency of the mutations in the heterogenous cell population. Measuring the frequency of the mutations would directly inform optimization of CRISPR procedures and amounts of effort necessary for downstream clonal isolation.

## Results

### SNP mutations in *mPer* genes are generated efficiently in mice by CRISPR-Cas9

Although key circadian parameters of the clock seem to be encoded in the PERIOD (PER) protein, it is little understood how 24 hr time cues are generated by the regulation of PER. As with most other proteins, PER has a modular structure with multiple domains, including the PAS domain, CRY-binding domain (CBD), and CK1-binding domain (CKBD) (Fig S1). Although the CBD and CKBD are known to be binding domains for other proteins (hence their names), their roles in generating 24 hr posttranslational oscillations of PER in phosphorylation and degradation remain poorly characterized. Although PAS is known as a homo or hetero-dimerization domain and conserved from plants to animals as an essential timing device^48, 49^, its role in mammalian clocks is also little studied. To understand how the PAS domain contributes to rhythm generation of PER at posttranslational level, we decided to disrupt the main function of PAS, homodimerization, in MEFs and mice.

Because the dimer structure of PER and PER PAS domains has been solved at high resolution by X-ray crystallography^50, 51^, we targeted critical motifs and amino acid (AA) residues for dimerization based on these available data. According to the structural studies, motifs containing PER1-W448 and PER2-W419 AA are most critical for dimerization (Fig S1). Tryptophan to glutamate mutations (PER1 W448E and PER2 W419E) have been suggested to be most disruptive to the hydrophobic interaction-mediated dimerization; we therefore planned to generate these mutations using CRISPR for functional studies.

Before we initiated the project in mouse models, feasibility of the project was tested in a clock cell model U2OS which has been proven to have a functional clock and to be amenable to CRISPR genome editing^17, 47, 52^. When the motifs harboring W448 in PER1 and W419 in PER2 were targeted by CRISPR, diverse indel mutations were generated including in-frame mutations leading to deletion of several AAs (Fig S2). Interestingly, all of the in-frame AA deletion mutants exhibited defective phosphorylation: largely truncated or hypo-phosphorylation of both PER proteins compared to wt PER. A subset of these PER deletion mutations as well as PAS domain point mutations W448E in *mPer1* and W419E in *mPer2* were tested in a different human cell line (HEK293) through transient transfection using mammalian expression plasmids; transient co-expression of CK1δ produced much less phosphorylation of these mutant PERs than wt PER (Fig S2). This is exciting but counterintuitive because PAS has not previously been directly implicated in PER phosphorylation or interaction with CK1.

All the data above strongly indicate that the PAS domain should be critical in circadian timing because PER phosphorylation is the basis of the mammalian circadian timer; thus mutations in and around the critical tryptophan residue that disrupt phosphorylation should impair circadian rhythms. As discussed above, we aimed to generate tryptophan-to-glutamate mutations in *mPer1* and *mPer2*. We selected an efficient sgRNA sequence close to the residues by testing several sgRNA sequences around the residues. MEFs were transfected with all-in-one pAdTrack-Cas9-sgRNA plasmid followed by FACS sorting for positive cells (GFP from the all-in-one plasmid)^17^. These cells were subjected to T7E1 assays and the most effective ones were selected (Fig 1a). When ssODNs for homology-directed repair (HDR) to mutate W448E in *mPer1* and W419E in *mPer2* were designed, novel restriction enzyme sites were added in the template to facilitate genotyping without sequencing PCR amplicons (Fig 1b). These additional mutations are also necessary when digital PCR is designed to distinguish between wt and mutant alleles. Although annealing temperature is different with single nucleotide polymorphism, it is very challenging to develop an all-or-none annealing condition based on a single nucleotide mutation^53^. CAS9 protein along with sgRNA and ssODN were injected into one-celled fertilized eggs to produce two knockin mutant mice. These injections generated 79 and 91 live pups for *mPer1* and *mPer2* mutant mice, respectively. As summarized in Fig 2a, we obtained 34 heterozygous (het) and 9 homozygous (ho) mutant mice for *mPer1*^*W448E*^. Based on the total number of alleles (158) we examined, KI efficiency was 33%. In addition, we obtained 13 useful in-frame AA indel mutant mice in which we expect a more severe phenotype than the point mutation. For *mPer2*^*W419E*^, 24 heterozygotes and no homozygotes were produced (Fig 2b). Although enzyme digestion of PCR amplicons suggested several homozygotes (see clones with red asterisk in Fig 2b), they turned out to be all heterozygotes when assayed by digital PCR (see below). One of two alleles had large deletions in these mice, which could not be amplified with the primer set and thus produced an unmixed sequencing chromatogram. We do not believe that fewer *mPer2* indel/mutant mice were produced because DSB was less efficiently made at *mPer2* W419 by the sgRNA. Injection for *mPer2* KI was done at three different times while the whole injection for *mPer1* KI was done at one time. The first batch of *mPer2* injection was most successful and the efficiency for indels/mutation in this batch was comparable to that of *mPer1* KI suggesting that the low efficiency could be due to failed delivery of CRISPR reagents into some of the fertilized eggs.

**Fig 1.**
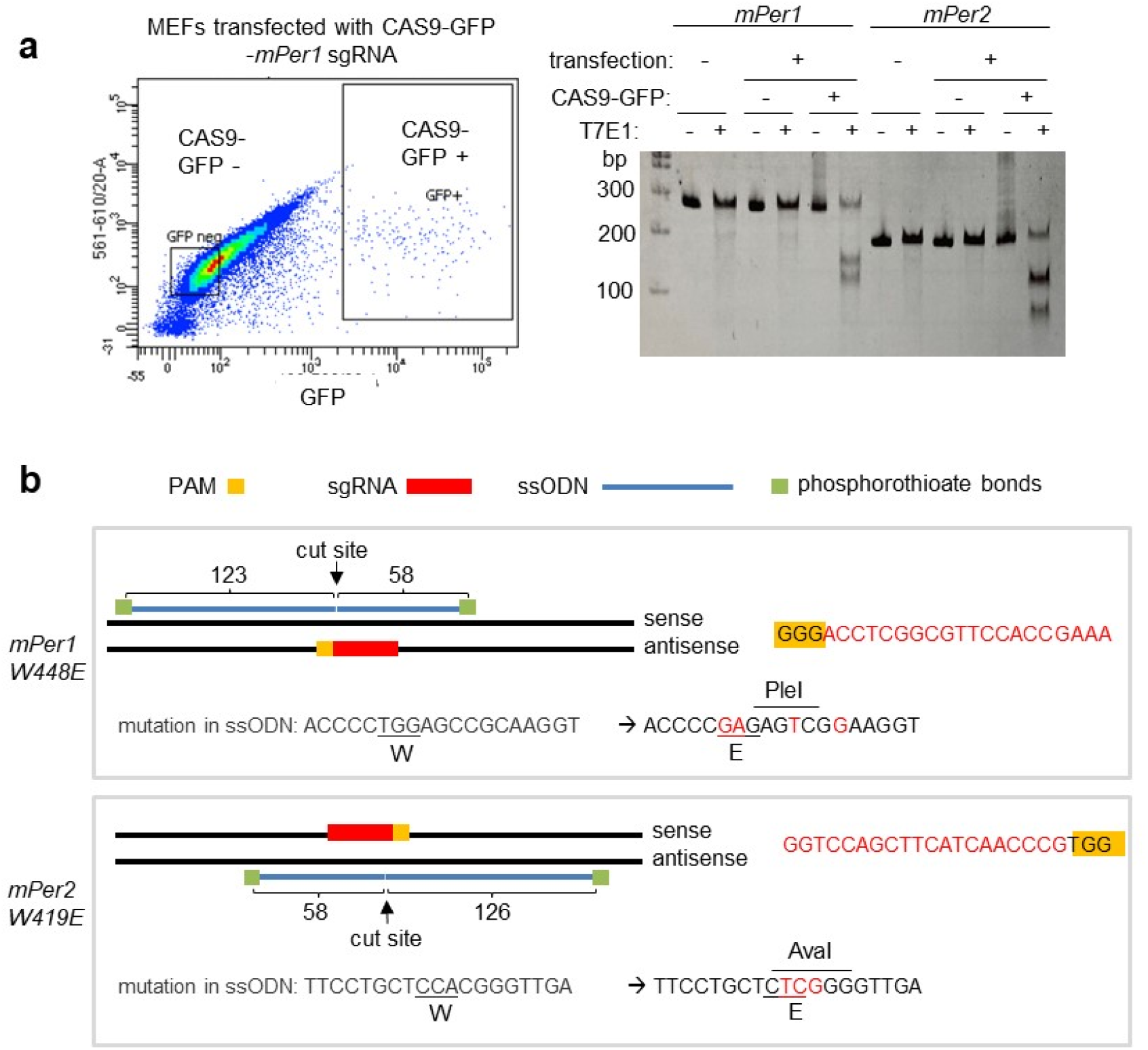
Strategy to knock-in mutations into *mPer1* and *mPer2* genes. **(a)** Efficient sgRNAs targeting *mPer1 W448* and *mPer2 W419* residues were selected by T7E1 assays in MEFs. Positive and negative transfected cells were selected by GFP expression and subjected to T7E1 assays. T7E1 results for less efficient sgRNAs are not shown. **(b)** ssODNs for HDR are designed based on location of sgRNA. Note the asymmetric homologous arms.

**Fig 2.**
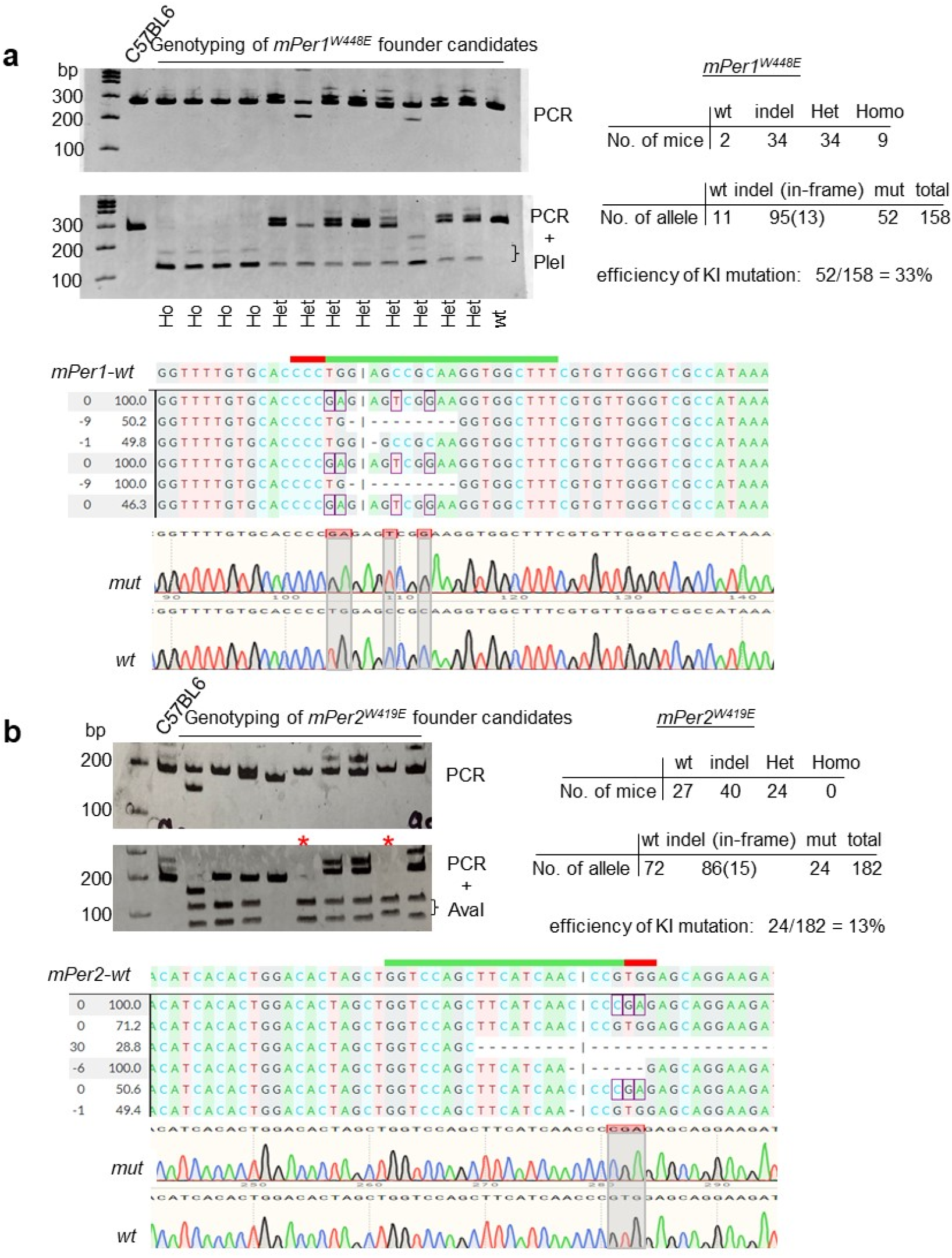
Knockin mutant mice are efficiently made by CRISPR-Cas9. **(a, b)** Genotyping by PCR and enzyme digestion showed efficient genome editing for two knockin mutations. Note that PCR amplicons indicated by red asterisk in (b) were completely digested by AvaI. Some of the indel mutants along with the specific knockin mutants are shown by Sanger sequencing.

### Genotyping for CRISPR-mutant mice can be streamlined by digital PCR

From the conventional genotyping by PCR and enzyme digestion (Fig 2), we believed 5 homozygotes for *mPer2*^*W419E*^ mutant mice were made because PCR amplicons from these mice were completely digested by AvaI. However, we did get only 50% instead of 100% heterozygous pups from breeding between these mutant and wt mice suggesting that one allele in these mice had large indels and thus the small PCR amplicon using the primer set could not be generated from these alleles. Indeed, PCR producing a large amplicon, ∼1.2 kb revealed large deletion alleles in three of the mice (Fig S3a). Despite several attempts using primers generating ∼ 3kb amplicons, we were not able to produce mutant PCR amplicons from the remaining two mice. Since large indels are not rare events and correct genotyping is critical when selecting founder mice, we developed a droplet digital PCR (ddPCR) assay for genotyping, which can quantify the number of the mutant copy over the copy number for a reference gene in the same sample in a PCR reaction and thus reveal correct genotypes regardless of indel size and digestion pattern. If a mutant probe is used, the ratio would be 1, 0.5 and 0 for homozygotes, heterozygotes and wt mice, respectively (Fig S3b).

Sensitivity and accuracy of ddPCR were assessed by generating standard curves for low copy numbers of the mutant genomic alleles spiked into 5,000 copies of the wt genomic allele (Fig 3a, 3b and S4). When 20, 100 and 500 copies of the mutant alleles were spiked into 5,000 copies of the wt allele, the ddPCR produced a strong linear correlation between added and detected copies (Fig 3a and b) (R^2^>0.98). The reference probe for *RPP30* gene also consistently detected ∼5,000 copies in these samples. Genotyping of both *mPer* mutant mice using this ddPCR protocol produced reliable results which were consistent with those obtained by enzyme digestion of PCR amplicons, except for five *mPer2*^*W419E*^ mutant mice (Fig 3c-f). Three of the *mPer2* mutant mice were compound heterozygotes, which matches the results of PCR genotyping producing a larger amplicon (Fig S3a). The other two compound heterozygous *mPer2*^*W419E*^ mutant mice with putative large deletions could be also confirmed by ddPCR (Fig 3d and f). ddPCR with the mutant probe on these samples produced ½ signals relative to those of the RPP30 probe whereas the wt probe did not produce any signal. These results demonstrate that the conventional PCR-based genotyping may not be suitable for all CRISPR mutant mice, especially ones with large indels because optimizing PCR conditions can be very challenging if not impossible.

**Fig 3.**
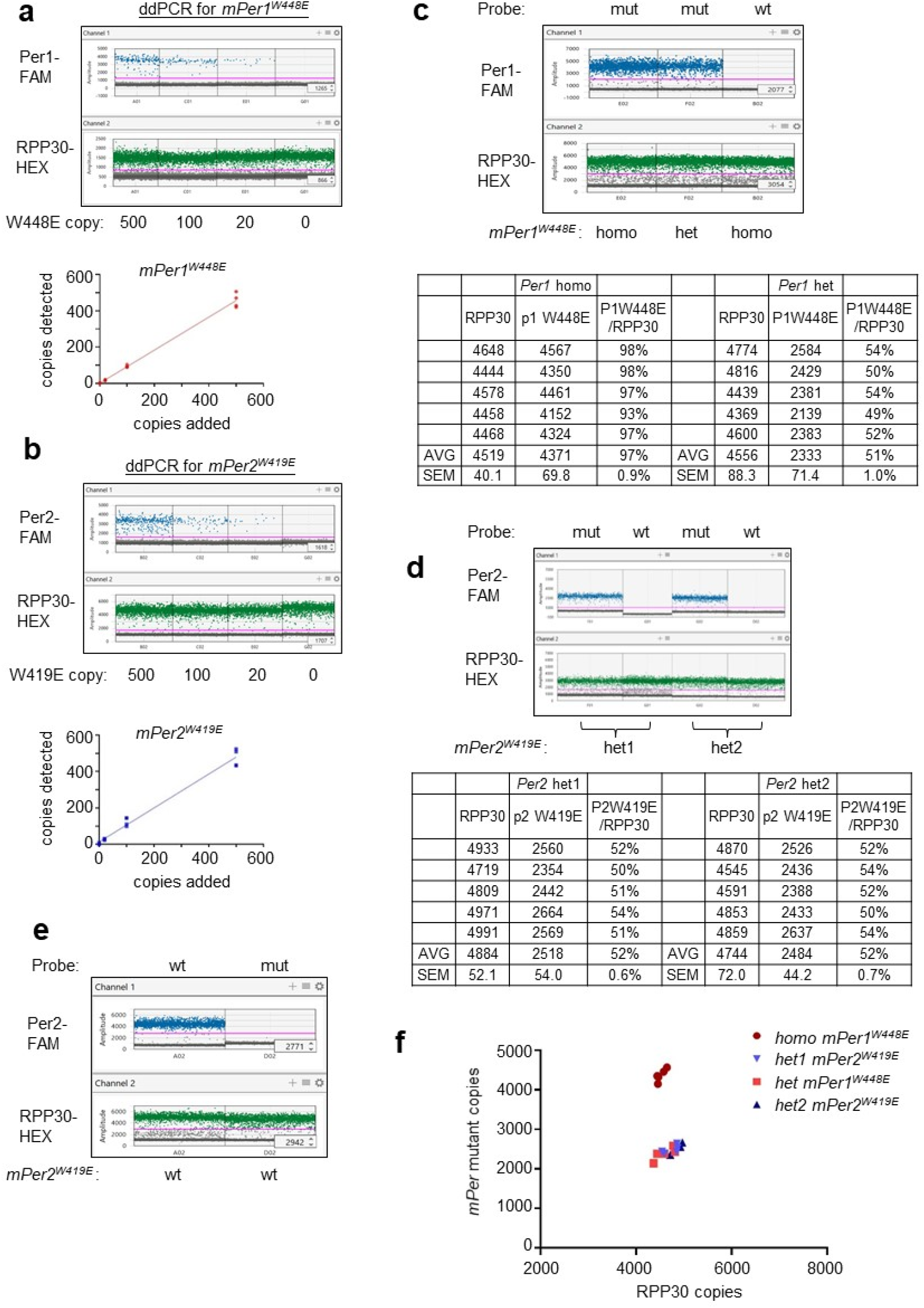
Digital PCR can be used to genotype CRISPR mutant mice that cannot be genotyped by PCR analysis. **(a, b)** ddPCR was optimized using genomic DNA obtained from the mutant mice in Fig 2. Our ddPCR condition can reliably detect as low as 8 mutant copies in a reaction when they are mixed with 5,000 copies of wt allele (Fig S4). N= 4 each. **(c, d)** ddPCR produced reliable genotyping results even in compound heterozygotes which could not be genotyped by PCR and enzyme digestion. PCR amplicons could not be generated from one of two alleles in two compound heterozygotes in (d) because they presumably had large deletions. Note that the wt probe could not detect wt allele in these heterozygotes. N= 5 each. **(e)** The wt probe can detect wt allele. **(f)** Digital PCR can be a streamlined procedure for rapid genotyping of CRISPR mutant mice even for mutant mice with large indels.

### Targeted mutations by CRISPR-Cas9 are significantly less efficient in MEFs compared to mice

Although CRISPR genome editing can be done in vivo in a much more efficient manner compared to the conventional knockin methods, it still requires a lot more resources including space for animals compared to cell-based systems. As with many biological fields, circadian mechanisms can be studied in cells because the circadian clock is cell autonomous. Although demand for genome editing in cells in the circadian field is increasing because molecular mechanisms can be better studied in cell models such as MEFs and U2OS, there are only few cases where specific mutations other than knockouts in clock genes have been made in these cells. Many clock mutant mice are readily available, but the polymorphisms in clock genes in the human gene pool are orders of magnitude larger. Implementing a streamlined workflow for specific genome editing such as introducing SNPs or short mutations in MEFs derived from existing mutant mice would greatly facilitate mechanistic studies of diverse genetic conditions. The approach will be also valuable in other fields where non-transformed cell models such as MEFs are required.

To streamline specific genome editing in MEFs, we assessed the efficiency of generating the *mPer1*^*W448E*^ and *mPer2*^*W19E*^ mutations using the same sgRNA and repair templates in MEFs that we had used for CRISPR editing of mouse zygotes. To compare the efficiency in a quantitative manner and assess the downstream clonal isolation workload, ddPCR was performed on FACS-sorted positive cells after transfection of the reagents as described in Fig 1a. In bulk MEFs transfected with the same sgRNA and ssODN for *mPer1*^*W448E*^ mutation, ∼3% of wt *mPer1* were converted into the mutant allele (Fig 4a). The efficiency was 5-6% for *mPer2*^*W419E*^ (Fig 4b). Although these numbers are significantly lower than those in mice, it is still very reasonable for researchers to successfully isolate desirable mutant clones by screening only a few dozen random clones using simple molecular techniques such as enzyme digestion of PCR amplicons as described above. To verify this efficiency in isolated MEF clones, some of the bulk positive cells above were singly sorted into 96 well plates and expanded for the PCR analysis followed by enzyme digestion. When we screened 15 clones each, 1 heterozygote clone for *mPer1*^*W448E*^ and 1 heterozygote clone and 1 homozygote clone for *mPer2*^*W419E*^ were isolated, roughly matching the ddPCR numbers (Fig 4c and d). The ddPCR assay is also a powerful tool when CRISPR-HDR protocols are being optimized. Efficiency of HDR was greatly affected by the ratio of CRISPR plasmid to donor template in co-transfection (Fig 4e).

**Fig 4.**
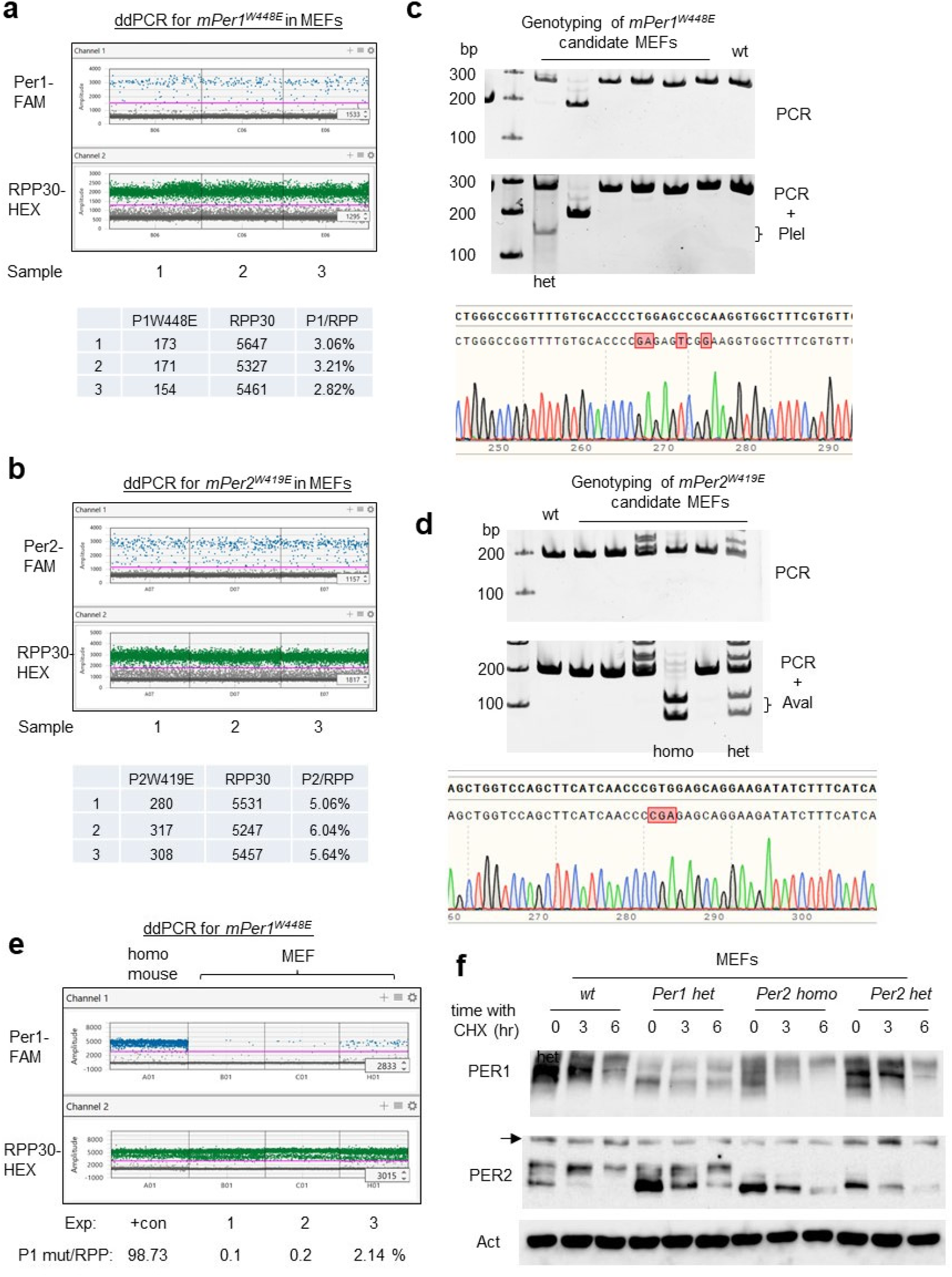
Digital PCR can be used for accurate quantification of specific genome editing in MEFs. **(a, b)** Frequency of two *mPer* mutations were quantified by ddPCR in MEFs transfected with the CRISPR reagents. ddPCR was performed on three independent samples each. **(c, d)** *mPer* mutant clones were successfully isolated from screening of 15 random clones each. Genomic DNA from these clones were amplified by PCR and digested with PleI (*mPer1*) and AvaI (*mPer2*). **(e)** Digital PCR can be a powerful tool to optimize CRISPR-HDR protocols. Varying amounts of *mPer1*-HDR template and all-in-one Cas9-sgRNA plasmid were co-transfected into MEFs followed by FACS sorting. Genomic DNA from a homozygous *mPer1* mutant mouse was used as a positive control. (f) PAS mutant mPER proteins show defective phosphorylation. Although both PER1 and PER2 mutants showed defective phosphorylation, *mPer2*^*W419E*^ mutant is apparently more defective in phosphorylation.

When PER proteins were probed in these mutant MEF clones, both PER proteins were dramatically less phosphorylated compared to control MEFs (Fig 4f). Existing PER proteins are progressively phosphorylated when do novo translation is inhibited by cycloheximide (CHX). These data along with the data in Fig S2 strongly suggest that PAS dimerization is critical for PER phosphorylation and probably for the clockwork as well. Because MEFs are not transformed cancer cells and thus are frequently used as an in vivo model, our data strongly support that MEFs can be an effective platform to study in vivo cell physiology, and this is even more compelling by already available numerous genetic mouse models.

## Discussion

As with most biological fields, our current understanding of the circadian clock in mammals has been largely established by reverse genetics in mice that have led to identification of a dozen essential clock genes to date^54^. Although the mechanistic insights from these mutant mouse models cannot be overstated, the traditional mutant mouse models based on gene targeting by homologous recombination required tremendous resources and a long period (18-24 months) even to assess if desired mutant models were made. In that sense, CRISPR revolutionized how mutant animal models are generated. It is faster (4-6 months) and much more affordable and efficient (Fig 5). In addition, as in our case, specific genome editing by CRISPR generated useful by-products such as knockouts and AA indels. Because these AA indels would disrupt the function of the motif more severely than the single AA mutation in the motif—and the larger the indel, the more severe the phenotype may be—these indel mutants along with the specific mutant animals would provide more complete insights into the function of the motif and the whole protein. As in our study, compound heterozygotes would be the most predominant genotype with successful injection of CRISPR reagents and sgRNA efficiency. ddPCR assays are very useful for accurate genotyping of these compound heterozygotes, especially alleles with large indels.

**Fig 5.**
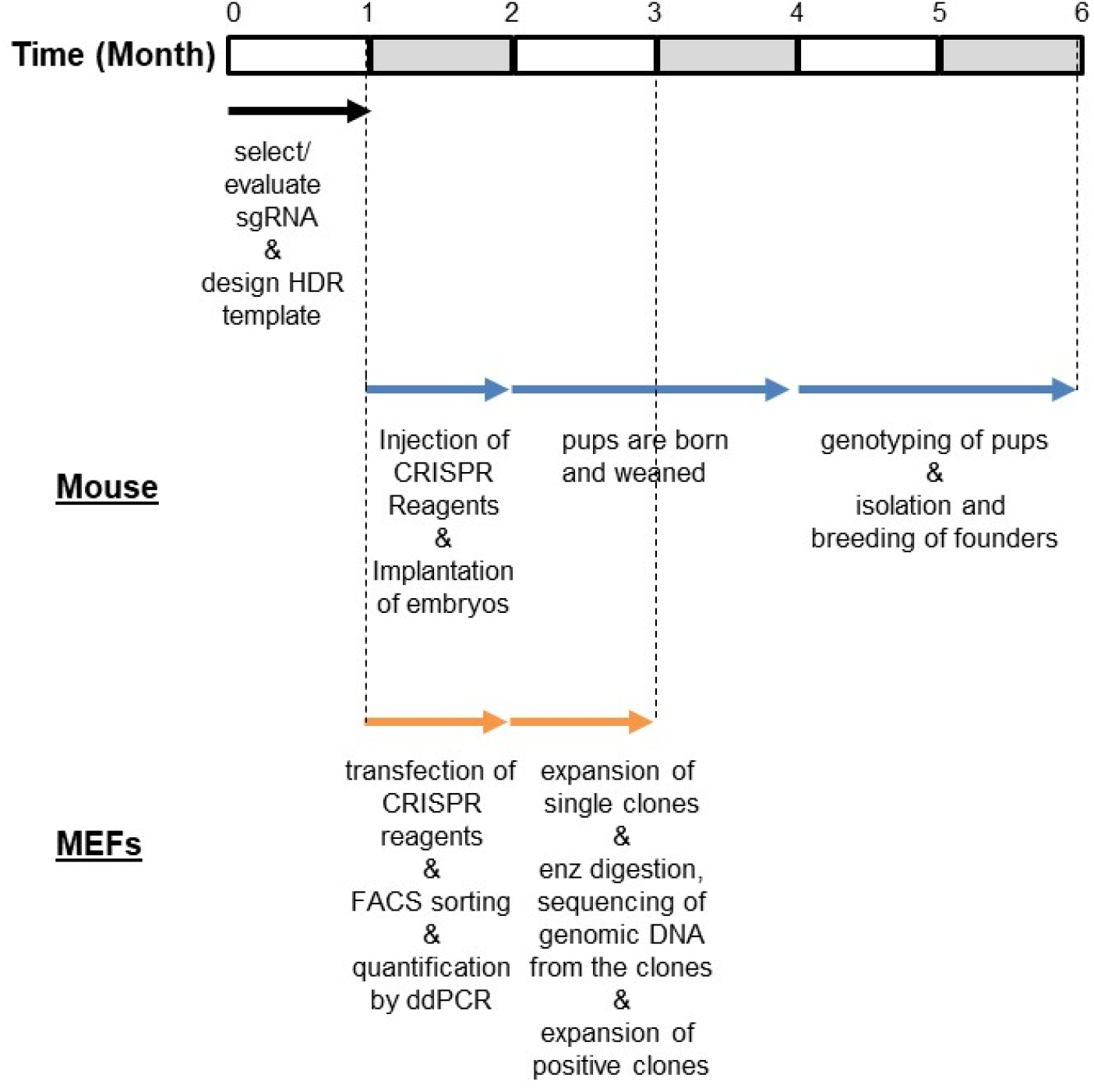
Timeline comparison in CRISPR-knockin mutation between mice and MEFs. Reasonable amounts of time for troubleshooting and optimization typical for a small laboratory are included in this timeline. From injection of CRISPR reagents into fertilized eggs to producing pups are usually done by a dedicated animal facility. The rest of the animal work and all the work in MEFs can be done by a laboratory.

Cell culture models will continue to be an effective platform for studying molecular mechanisms in many biological pathways including circadian biology, given their time- and cost-efficiencies. Although the genome in cell culture models can be precisely edited using CRISPR-Cas9, it is also important to recognize such genome edited cell models are very rare in general. This may reflect that precise genome editing is still not practical in many small laboratories because HDR frequency is significantly low compared to that of NHEJ, and thus it is challenging to isolate HDR clones out of much more abundant NHEJ mutant clones. Frequency of the specific mutation could be easily estimated by enzyme digestion followed by gel electrophoresis if a novel digestion site is added into the repair template, and the HDR frequency is high enough. However, we could not detect digested fragments from PCR amplicons prepared from genomic DNA of our bulk sorted MEFs on agarose gels. When HDR efficiency is 3-6% as in our bulk sorted cells, we believe the densitometric method is not sensitive enough to detect digested fragments on agarose gels. Next-generation sequencing (NGS) would be the holy grail for accurate quantification of HDR, but it is not practical especially when many mutant cell lines are generated, or HDR conditions are not yet optimized. Downstream work flow and effort would be greatly affected by the efficiency of HDR. For example, if the efficiency is ∼0.1%, at least several hundreds of random clones need to be expanded and interrogated by PCR analysis and sequencing, which would not be practical. Digital PCR assays can be a powerful tool when a specific mutant cell clone needs to be isolated, because frequency of the mutation can be accurately measured from a heterogenous population of mutant clones, which will allow researchers to estimate the number of single clones to be analyzed to isolate a few desirable mutant clones or to optimize the conditions to increase HDR efficiency (Fig 5).

As our understanding of many important biological pathways is becoming mature by identification of most of the essential genes, a next frontier in biology would be to learn how polymorphisms in these genes affect the pathways and physiology. It has been already demonstrated that some of these polymorphisms are associated with diseases like the circadian disorder called Familial Advanced Sleep Phase Syndrome^55, 56^. We believe our CRISPR method combined with ddPCR can be a streamlined process to generate any simple mutant KIs in MEFs to study human genetics, if relevant human physiology can be studied in MEFs like circadian biology.

Although we cannot generalize CRISPR efficiency for HDR in mice, available data suggest it is close to 100% if the sgRNA is properly selected and reagents are successfully delivered. Considering this high efficiency, the short time frame from CRISPR design to obtaining founder mice, and the potential to generate combined mutations by breeding, it may be more practical to generate mutant mice rather than individual mutant cell clones in some cases. Mouse models are obviously more versatile since whole animal physiology can be studied in single and combined mutant mice as well as molecular mechanisms in diverse cells or tissues isolated from these mutant mice.

In summary, when same sgRNA and HDR-template are used, HDR efficiency is dramatically higher in fertilized eggs compared to MEFs, and digital PCR is an extremely powerful tool for genotyping of these CRISPR mutant mice and for quantitative analysis of the mutation frequency if the genome editing is done in cells. Using our protocols, we believe that any simple mutant model can be efficiently generated in MEFs or other CRISPR-compatible cell lines, even in small laboratories because implementation of ddPCR is fairly straightforward.

## Author Contributions

K.L and C.L. planned and performed the experiments. K.L. and C.L. wrote the manuscript.

## Acknowledgments

We thank Dennis Chang for assistance with manuscript preparation. We thank Dr. Joseph Takahashi for arranging to use the transgenic core facility at UT Southwestern Medical Center. This work was supported by NIH R01 GM131283 (C.L.).

The authors declare no competing interests.

Correspondence and requests for materials should be addressed to C.L.

## Data availability

All data generated or analysed during this study are included in this published article (and its Supplementary Information files).

---

**Fig S1.**
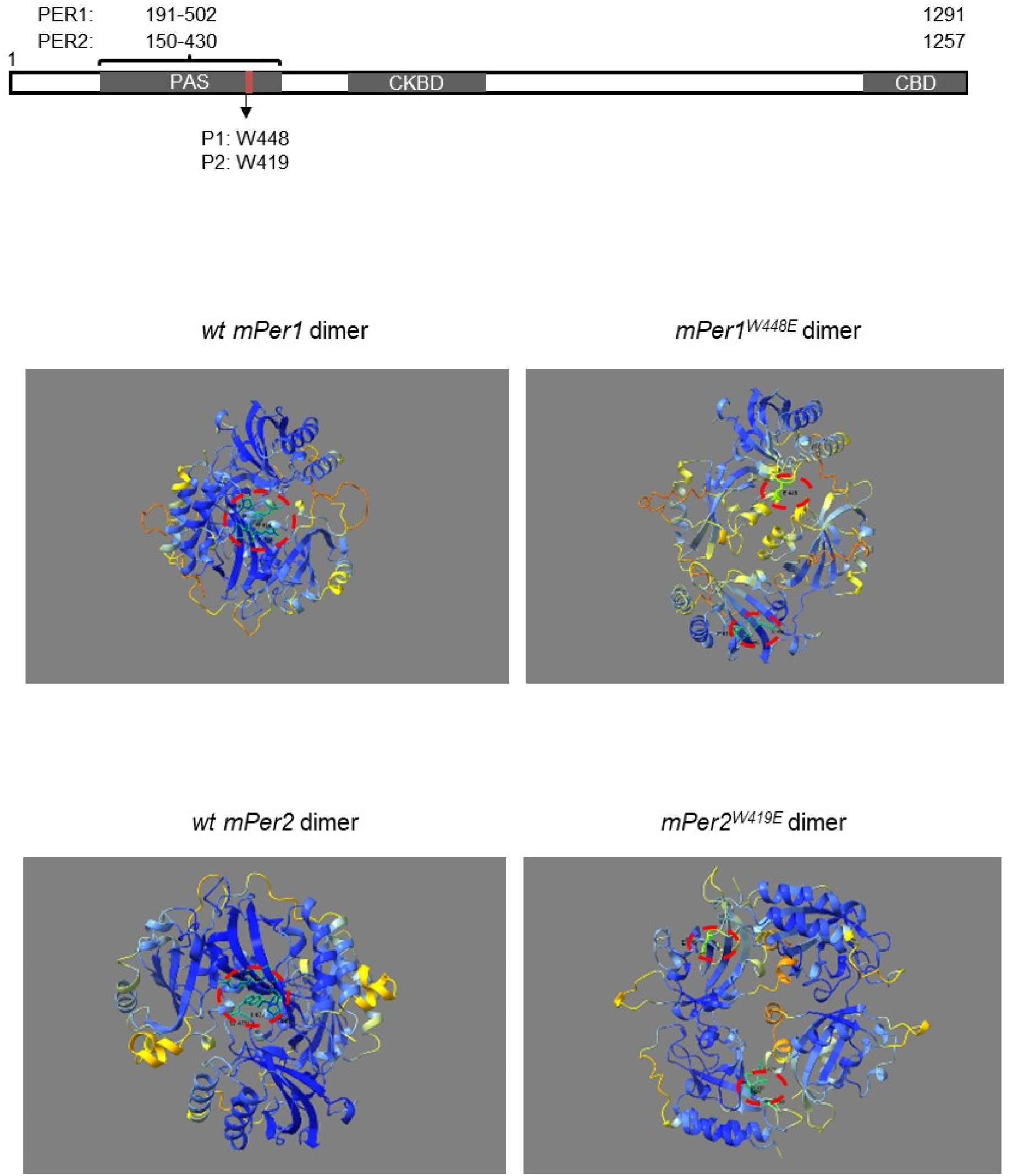
Mutations *mPer1*^*W448E*^ and *mPer2*^*W419E*^ are most disruptive SNPs in mPER1 and mPER2 homodimerization. When the mutations were introduced into PAS dimers and simulated by the AlphaFold program, the hydrophobic bonding indicated by red circles was completely disrupted. Mutant PAS homodimers are simulated based on the published wt PAS dimer structure.

**Fig S2.**
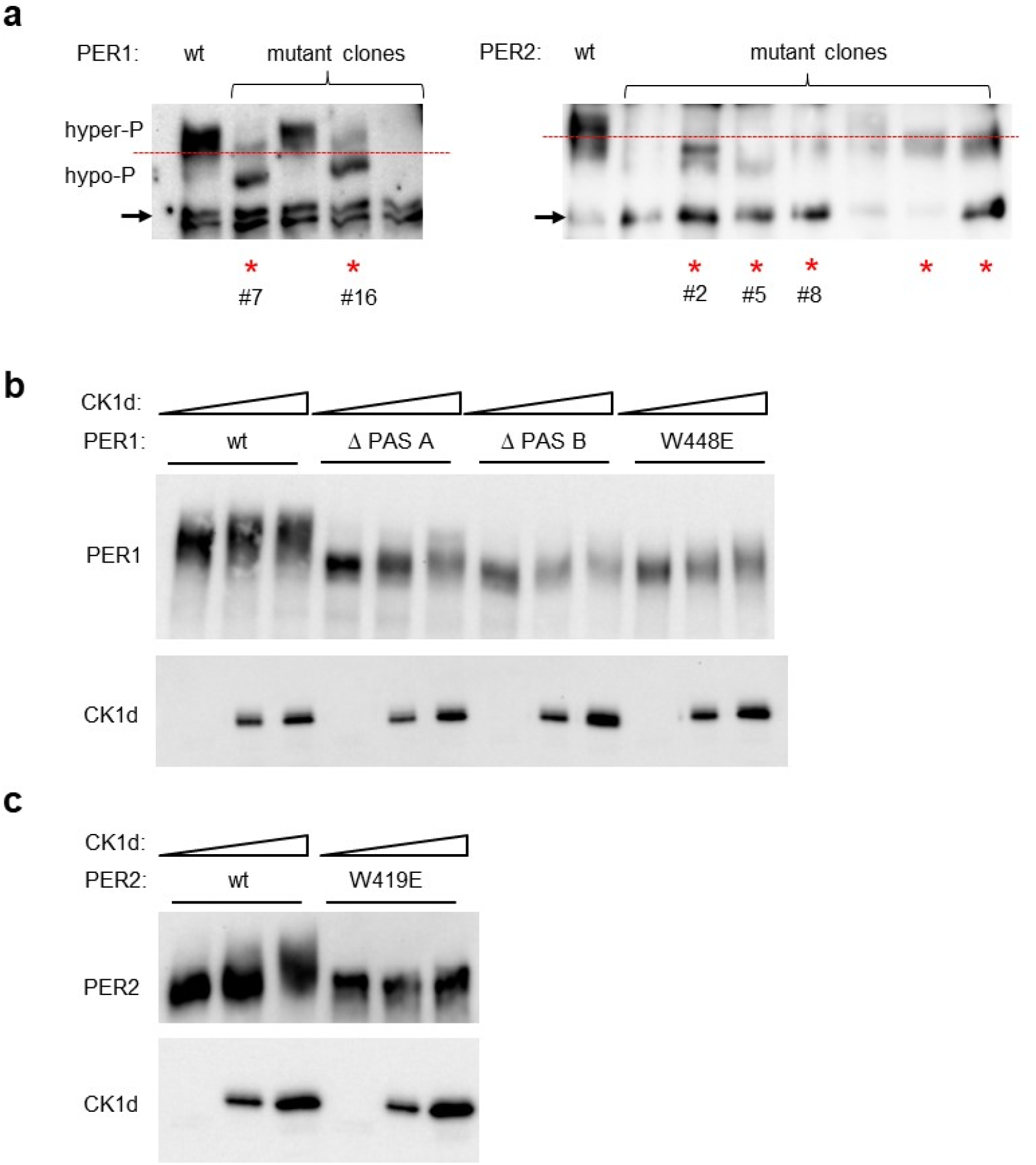
Random AA deletions in PAS B domain induce defective phosphorylation in U2OS cells. **(a)** hPER1 W448 and hPER2 W421 (conserved residues in human PER proteins) were targeted by CRISPR without HDR templates inU2OS cells. Note that PER proteins indicated by red asterisk show all hypophosphorylated PER compared to wt PER. The mutant *Per* genes from the numbered clones were sequenced (see the materials and methods). **(b, c)** When mutant mPER proteins with defective PAS domains were transiently co-expressed with CK1δ in HEK293 cells, they were all hypophosphorylated similar to the mutant endogenous hPER in U2OS cells. Δ PAS A and Δ PAS B represent mutant PER1 with deletions of AA 244-252 in PAS A and AA 444-448 in PAS B domain, respectively. Three different amounts of CK1δ were used: 0, 5 and 25 ng. *mPer* was fixed at 300 ng.

**Fig S3.**
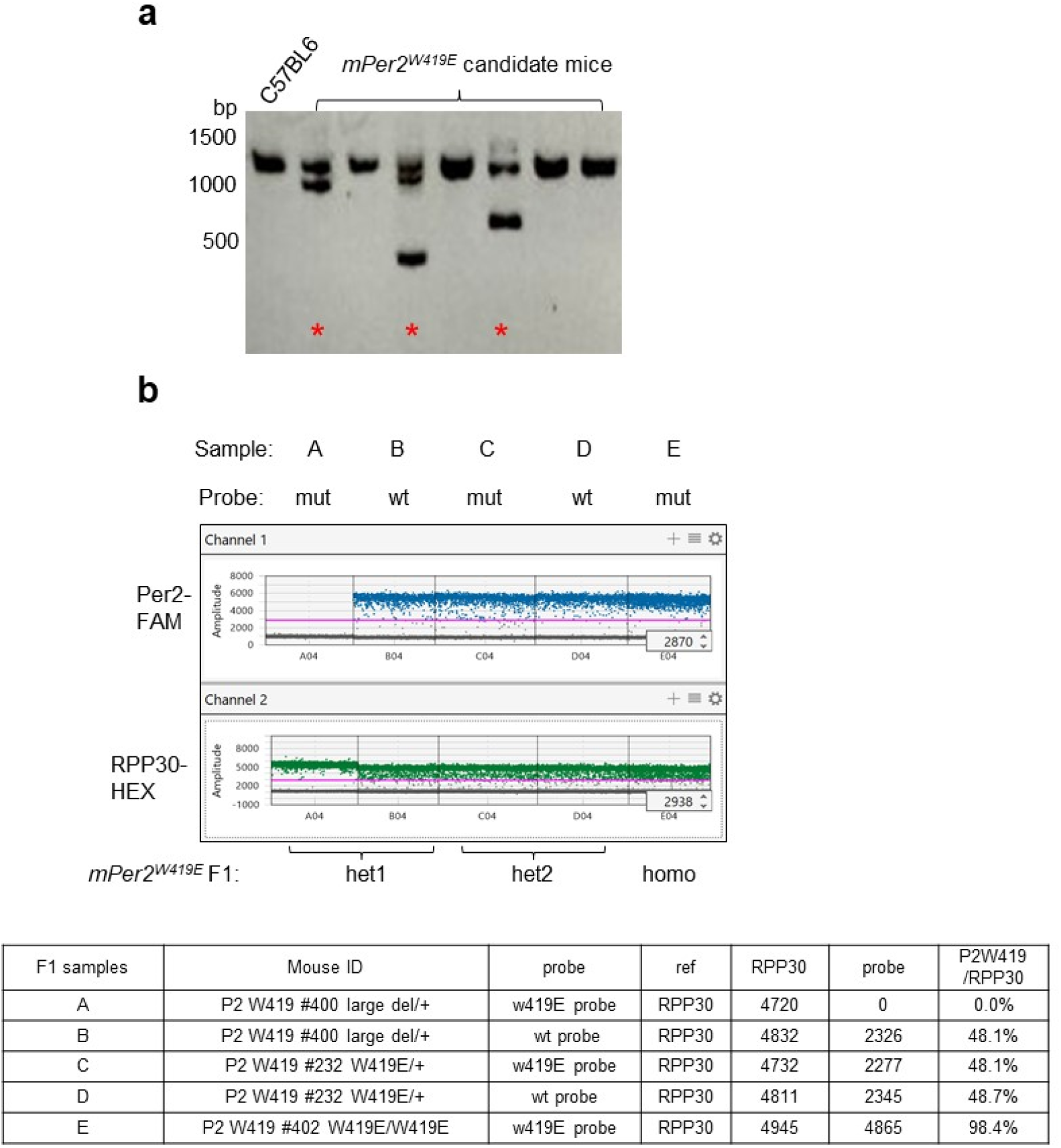
Digital PCR can be used to genotype CRISPR mutant mice with large indel alleles which cannot be analyzed by conventional PCR. **(a)** Increased PCR amplicon size can detect some large indel alleles. The amplicons indicated by red asterisks show that deletions in one allele in these mutant mice are larger than the amplicon size in Fig 2. (b) Digital PCR genotyping of F1 pups between wt and apparent mPer2 homozygotes revealed that the PCR analysis produced incorrect genotyping results of the founder mice.

**Fig S4.**
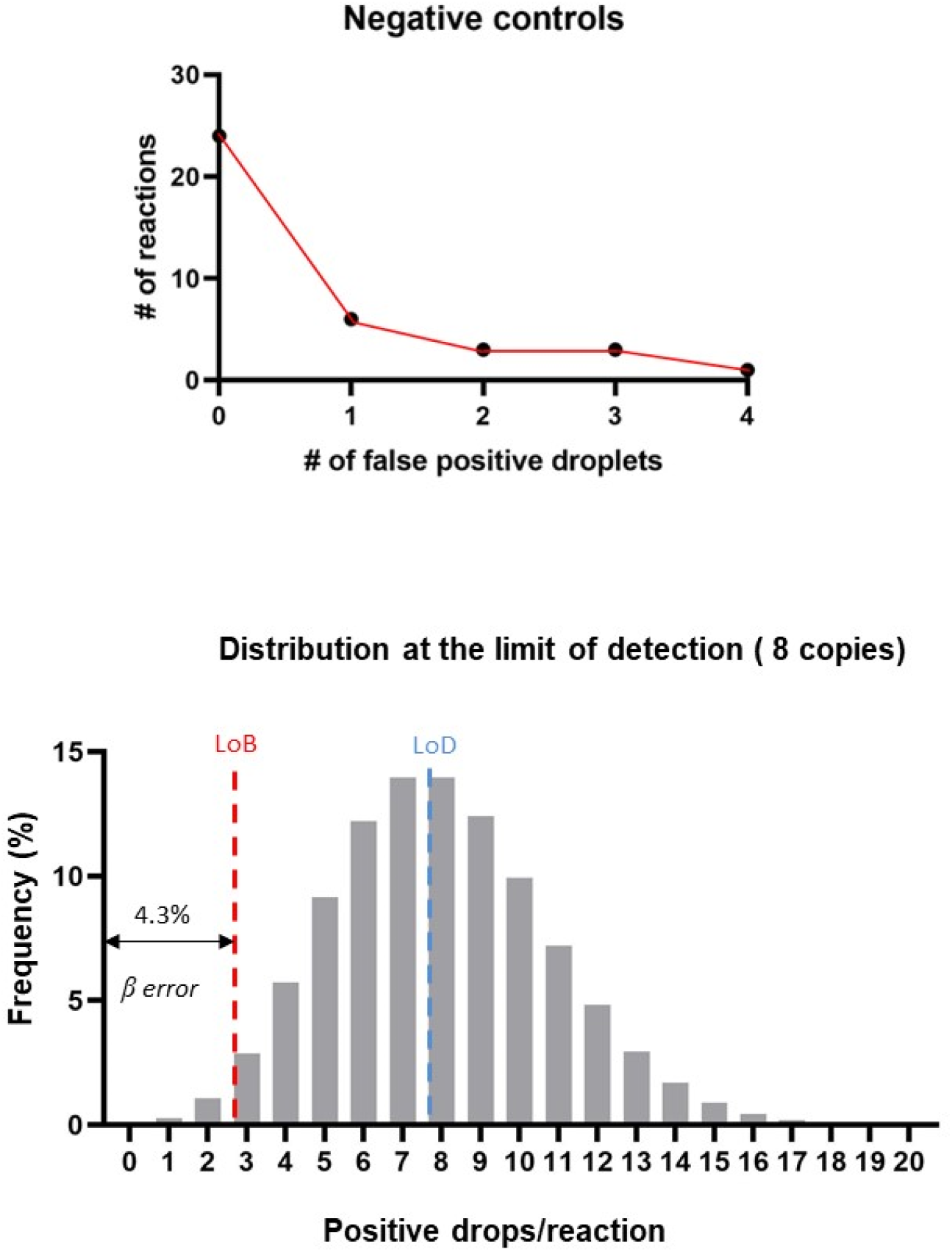
Our ddPCR condition can detect 8 copies of the mutant alleles in heterogenous MEFs. The limits and confidence intervals of our assays were established by determining a limit of blank (LoB) and a lower limit of detection (LoD) using our serial dilution samples. There were 24 positive drops in total 38 reactions giving an average false positive rate (Afp) of 0.71, LoB of 3 drops/reaction with α error<5%. The LoD was determined by a minimum droplet number with β error<5%, which was achieved at 8 drops/reaction. These numbers were not affected by total genomic DNA copy numbers in the reaction. The bottom graph represents the theoretical Poisson distribution when a sample is prepared at the LoD (8 copies/reaction). The arrow to the left of the LoB is β error.

## Materials and Methods

### Generation of mutant mice and genotyping

All mice were maintained in a climate-controlled room and used according to the Florida State University Animal Use Committee’s guidelines. All experiments involving animals were performed according to approved protocols. We used about equal numbers of male and female mice.

sgRNA and ssODNS were ordered from IDT. Injection of ribonucleoprotein complex consisting of Cas9 protein and sgRNA along with ssODNs into one-day fertilized eggs, and generation of pups from pseudo pregnant mice were done at the UT Southwestern medical center core facility. Tail tissues were obtained when these mice were weaned at 3 weeks old. PCR amplicons were obtained using the following primers, digested with PleI for *mPer1*genotyping and AvaI for *mPer2* genotyping and sequenced.

*mPer1_W448* forward primer: TGACCATTCCCCTATTCGCTT

*mPer1_W448* reverse primer: CCATGCCATGTCCATACCAC

*mPer2_W419* forward primer: TTCGATTATTCTCCCATTCGAT

*mPer2_W419* reverse primer: GAGAGGTGAGAATAGGCCAAAA

In the majority of the clones, two different Sanger sequencing traces were mixed due to different indels and/or the HDR mutation in two alleles. These results were deconvoluted by a computer algorithm called DECORD v3 (https://decodr.org/) into two separate traces. Accuracy of the deconvolution was confirmed by TA cloning if the decoding results were ambiguous.

For Fig S3a, the following primers were used to generate ∼1.2 kb amplicon.

*mPer2_W419* forward primer: GATCTGATCGAGACGCCTGTG

*mPer2_W419* reverse primer: ATGGCTGCAACACAGACGAT

### Amino acid sequence of mutant *Per1* and *Per2* clones in U2OS cells

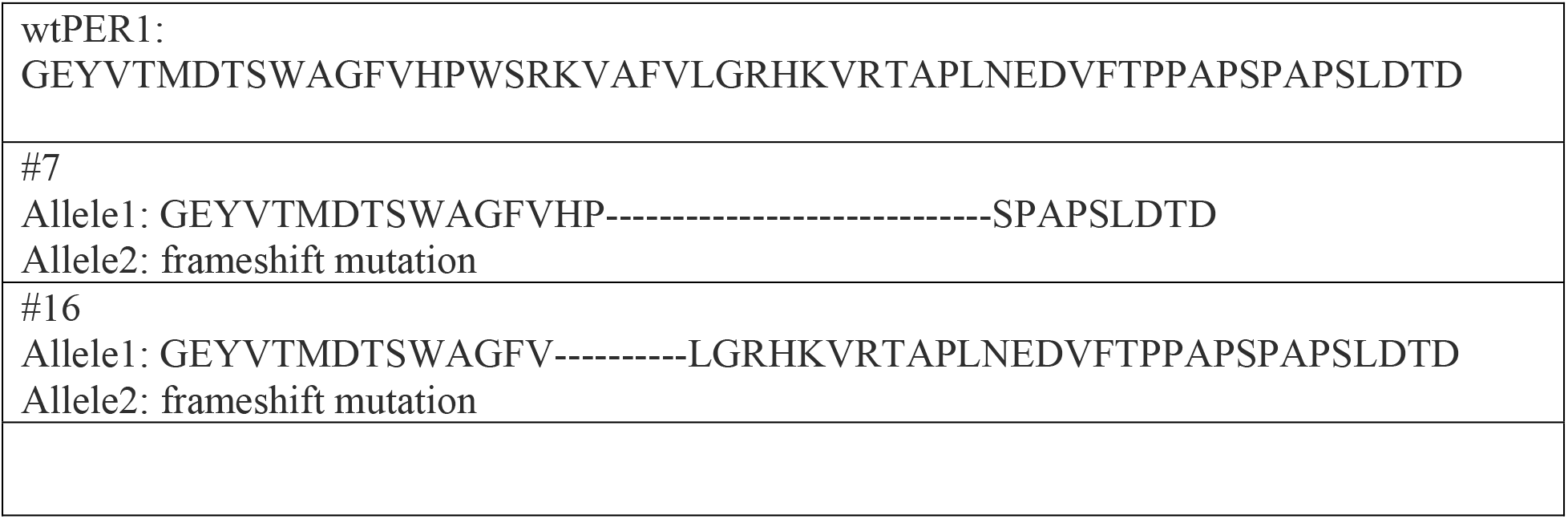
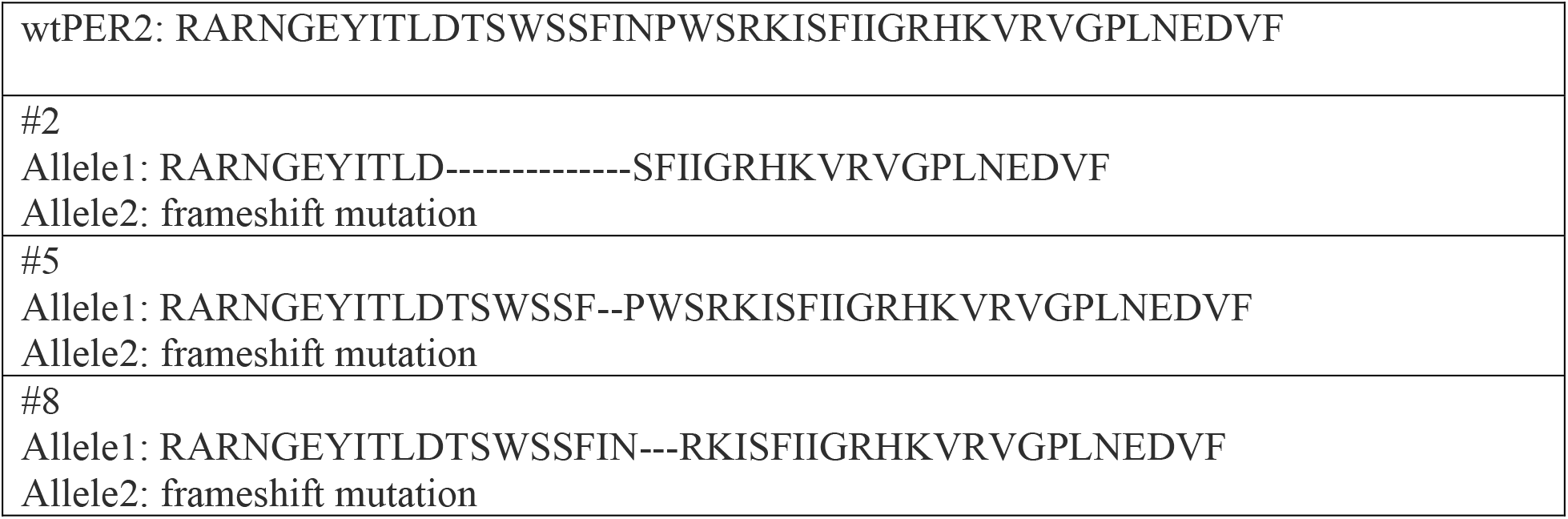

### Plasmid construction

*mPer1*^*ΔPAS A(244-252)*^, *mPer1*^*ΔPAS B(444-448)*^, *mPer1*^*W448E*^, and *mPer2*^*W419E*^ (Fig S2) were generated using Q5 Site-Directed Mutagenesis Kit (NEB E0554S). *pcDNA-mPer1*and m*Per2* plasmids were used as templates and have been described previously^37^. The pcDNA-CK1δ construct has been described previously^37^.

pAdTrack all-in-one vectors expressing GFP, Cas9 and sgRNA for human *Per* genes in U2OS cells were prepared by adding the following sgRNA sequence as described previously^17^.

sgRNA for *hPer1*: ggagccgcaaggtagccttc

sgRNA for *hPer2*: ggtccagcttcatcaaccca

### ssODN sequence

*mPer1*: CctgcgcccacagTACTGCAGCTGGCAGGCCAGCCCTTTGACCATTCCCCTATTCGCTTCTGTGCTCGGAACGGGGAATATGTCACCATGGACACCAGCTGGGCCGGTTTTGTGCACCC C <u>gaGAGtCGgAAGGTGGCTTT</u> CGTGTTGGGTCGCCATAAAGTGCGCACgtaagggaactgtg

*mPer2*: ttgccccctgctgtgagaggtgagaataggccaaaatcccccaaaacccacagagtggaaccctgggagcactcacACCCTGACTTTGTGCCTCCCAATGATGAAAGATATCTTCCTGCTCtcgGGGTTGATGAAGCTGGACCAGCTAGTGTCCAGTGTGATGTACTCCCCGTTGCGGGTGCGG

### Transfection, immunoblotting and T7E1 assay

For transfection with the all-in-one plasmids in U2OS cells (ATCC #HTB-96), cells were plated into 6 cm dishes to be approximately 60% confluent on the day of transfection. Cells were transfected with jetOPTIMUS according to the manufacturer’s protocol (jetOPTIMUS transfection reagent, Polyplus) and incubated for 2 days before they were subjected to trypsinization and FACS sorting using BD FACSAria SORP equipped with an Automated Cell Deposition Unit (ACDU) for sorting into 96 well plates. FACS-sorted cells were collected into two groups: negative and bright GFP, and plated into 35 mm dishes and grown for 2 days before harvest and genomic DNA extraction. Harvested cell pellets were homogenized in 250 ul solution A containing 0.1 M Tris-HCl pH=9.0, 0.1 M EDTA, and 1% SDS, and incubated at 70 °C for 30 minutes. 35 ul 8 M potassium acetate was added and incubated at room temperature for 5 minutes. The samples were centrifuged at 13,000 rpm for 15 minutes and genomic DNA was purified by subjecting the supernatant to phenol-chloroform extraction and ethanol precipitation. The extracted genomic DNA pellet was dissolved in 100 ul water and used as a template to PCR amplify the target genomic locus with a set of primers (Table 2) flanking the target region. Amplicons were confirmed by agarose electrophoresis, and purified with a PCR purification kit (QIAGEN Cat.28104). 200 ng DNA samples were denatured and annealed according to the protocol in Ran et al. ^4^ and subjected to T7E1 digestion in a 20 ul reaction according to the manufacturer’s protocol (NEB cat.M0302). Digestion products were resolved on 8% acrylamide/bis TBE gel and visualized with EtBr.

For transfection in MEFs, 900 ng of all-in-one plasmids were used for Fig 1, and 900 ng all-in-one plasmids + 300 ng repair templates were used for HDR mutations. For Fig 4e, 300 + 900 (1), 600 + 600 (2) and 900 + 300 (3) ng DNA were used for varying amounts of all-in-one plasmid to repair template, respectively.

For immunoblotting of U2OS and MEF mutant clones, single cells in 96 well plates were expanded and transferred to 35 mm and then 60 mm dishes. When the cells were confluent in 60 mm dishes, half of the cells in each dish were frozen and stored in a liquid nitrogen dewar and the other half were plated in one 60 mm dish for immunoblotting and one 35 mm dish for Sanger sequencing. Immunoblotting was done as described previously. Antibodies to clock proteins were generated as previously reported ^57, 58^. anti-PER1 (GP62), anti-PER2 (hP2-GP49) and CK1δ-GP antibodies were used at 1:1,000 dilution in 5% milk–TBST solution. Rabbit anti-ACTIN antibody (Sigma, A5060) was used at 1:2,000. For immunoblotting in Fig S2b a and c, expression plasmids were transfected into HEK293 cells.

### Droplet digital PCR (ddPCR)

Droplet digital PCR reagents were purchased from Bio-Rad and reactions were set up as follows.

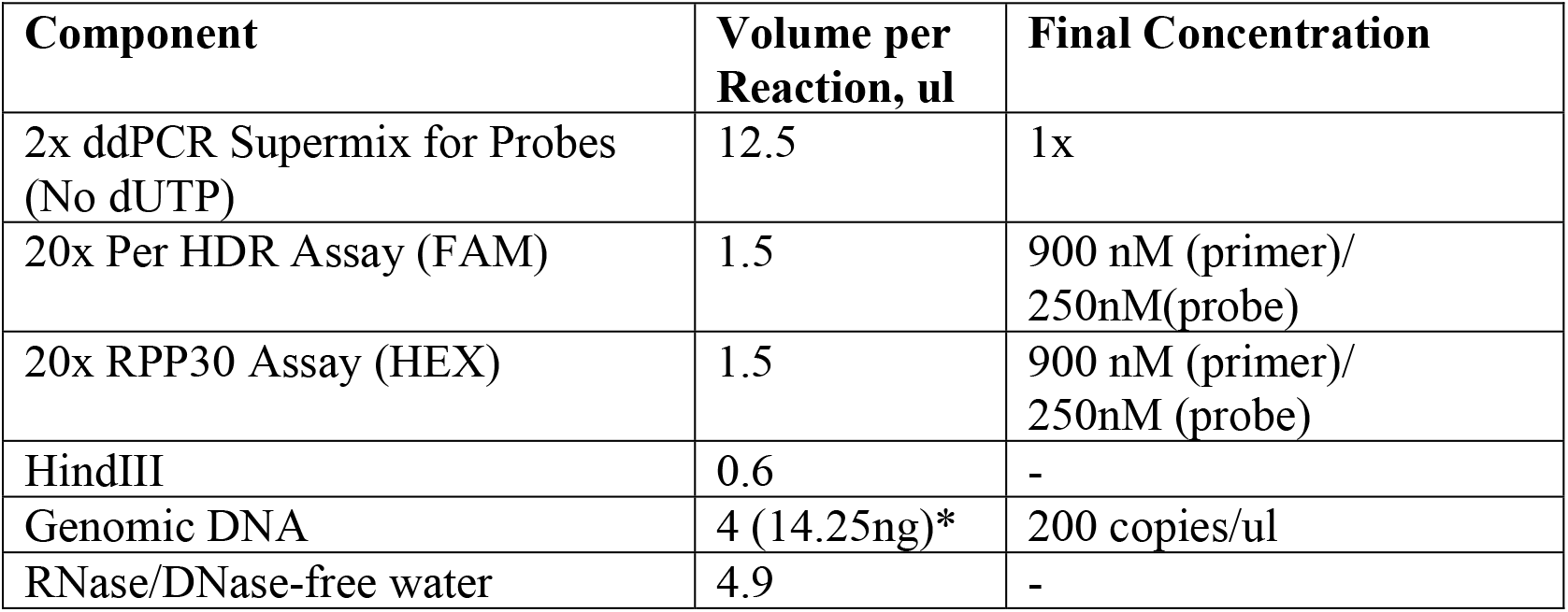
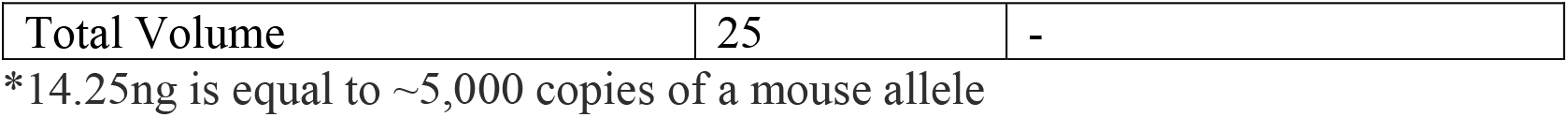

The following primers and probes were used.

ddPCR assay primers

*mPer1_W448* ddPCR forward primer: TGACCATTCCCCTATTCGCTT

*mPer1_W448* ddPCR reverse primer: CCATGCCATGTCCATACCAC

*mPer2_W419* ddPCR forward primer: TTCGATTATTCTCCCATTCGAT

*mPer2_W419* ddPCR reverse primer: GAGAGGTGAGAATAGGCCAAAA

*mRpp30* ddPCR forward primer: AAGAAACCACGGCCATCAGAAG

*mRpp30* ddPCR reverse primer: GGGTTTTATTTGCTGTTTTAATGGTC

ddPCR probes

*mPer1_W448* ddPCR wild-type probe:

ACCCCTGGAGCCGCAAGGT (/56-FAM/ACCCCTGGA/ZEN/GCCGCAAGGT/3IABkFQ/)

*mPer1_W448E* ddPCR mutant probe:

ACCCCGAGAGTCGGAAGGT (/56-FAM/ACCCCGAGA/ZEN/GTCGGAAGGT/3IABkFQ/)

*mPer2_W419* ddPCR wild-type probe:

TTCCTGCTCCACGGGTTGA (/56-FAM/TTCCTGCTC/ZEN/CACGGGTTGA/3IABkFQ/)

*mPer2_W419E* ddPCR mutant probe:

AGCTTCATCAACCCCGAGAG (/56- FAM/AGCTTCATC/ZEN/AACCCCGAGAG/3IABkFQ/)

*mRpp30* reference probe:

CTGCCTCCTCCCCTTCGTAG (/5HEX/CTGCCTCCT/ZEN/CCCCTTCGTAG/3IABkFQ/)

Droplets were generated from 20 ul of the samples and subjected to thermal cycles as follows.

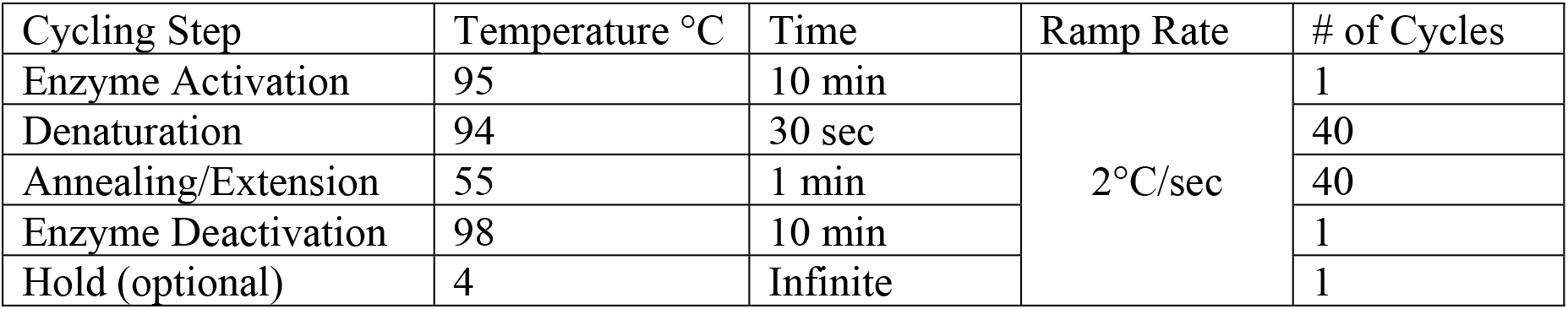

After the PCR amplification, the plate was transferred into the Bio-Rad QX-100 Droplet Reader. All assays were analyzed using the QX200 droplet reader and Quantasoft analysis pro (Bio-Rad).

### MEF single cell isolation and expansion, bioluminescence monitoring and immunoblotting

Immortalized *mPer2*^*Luc*^ MEFs were transfected and GFP-expressing cells were singly sorted into 96 well plates by FACS and expanded serially into 24 well plates, 6 well plates, and then 6 cm dishes as described above. Positive clones were isolated by enzyme digestion of PCR amplicons and Sanger sequencing and subjected to immunoblotting as described above.

### LoB and LoD of ddPCR assays

An average false positive rate (A_FP_) was measured from 38 negative samples. A total of 24 false-positive droplets was detected in 38 reactions (A_FP_ = 0.71). 97.4% of the reactions (37/38) contained <4 false positive droplets, while 89.5% of the reactions (33/37) contained <3 false positive droplets. The LoB was calculated and set at 3 to maintain an α error lower than 5%. α error indicates that the chance of detected false positive droplet counts in a negative control sample is greater than LoB. The LoD was calculated and set at 8 to maintain a β error lower than 5%. β error indicates that the probability of detected droplet counts is lower than LoB when a sample is prepared at the LoD ^59^.

### Prediction of protein folding

Protein folding of the wt vs. mutant dimers, mPER1 PAS (PDB: |4DJ2|191-502 vs mPER1[197– 502] W448E mutant) and mPER2 PAS (PDB: |3GDI|170-473 vs mPER2[170–473] W419E mutant) was predicted using ColabFold, a user-friendly version of AlphaFold^60,61^. Sequence coverage plots and a total of 5 models were predicted and ranked. Predicted Local Distance Difference Test (pLDDT) and predicted aligned error (PAE) were computed to measure the confidence of each predicted model. The best-ranked model was presented (Fig S1). The 3D structures were created, and the dimerization interface was visualized with UCSF ChimeraX ^62^.

