## Supplemental Figs for "Quantitative comparison of CRISPR-Cas9-mediated mutation efficiency between mice and MEFs using digital PCR assays"

Fig S1

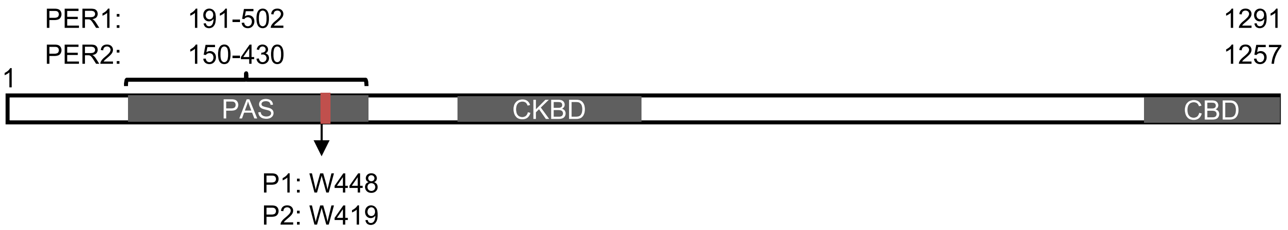


*wt mPer1* dimer

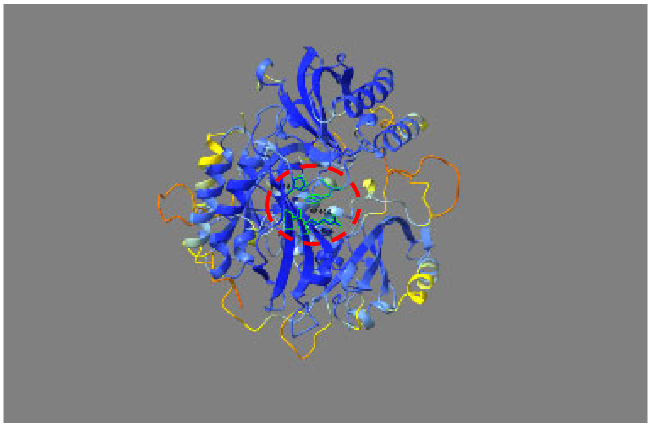

*mPer1$^{W448E}$* dimer

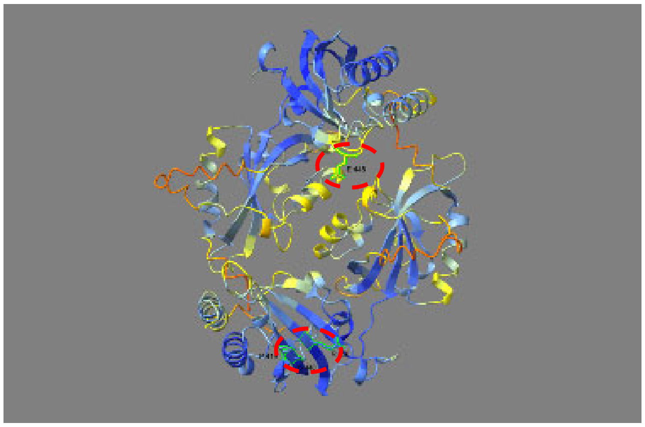

*wt mPer2* dimer

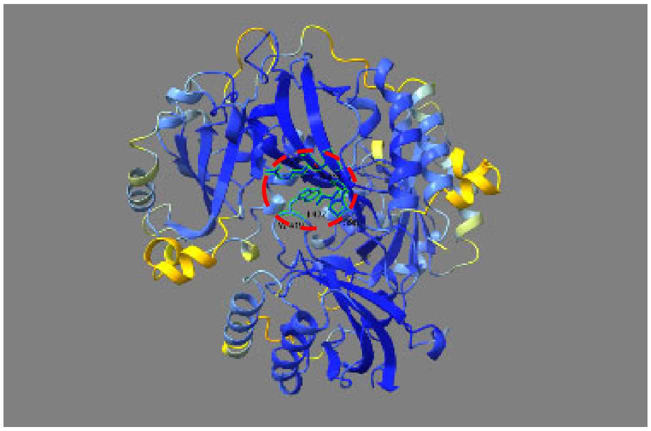

*mPer2$^{W419E}$* dimer

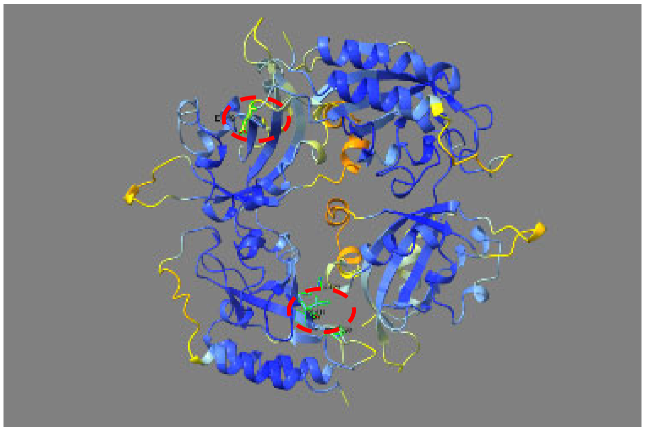

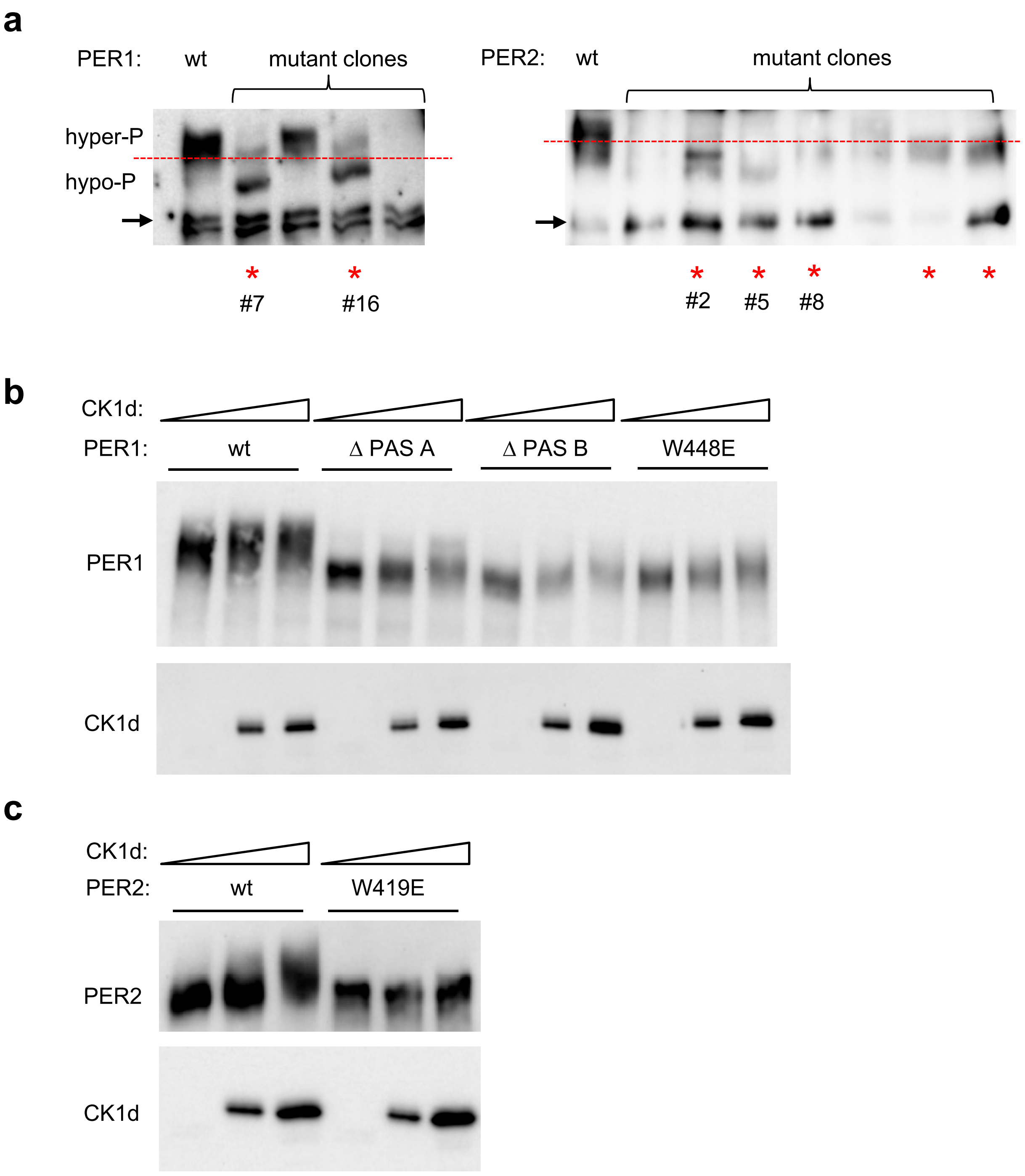
a
PER1:
wt
mutant clones
hyper-P
hypo-P
#7
#16
PER2:
wt
mutant clones
#2
#5
#8
b
CK1d:
PER1:
wt
Δ PAS A
Δ PAS B
W448E
PER1
CK1d
c
CK1d:
PER2:
wt
W419E
PER2
CK1d

Fig S3

**a**

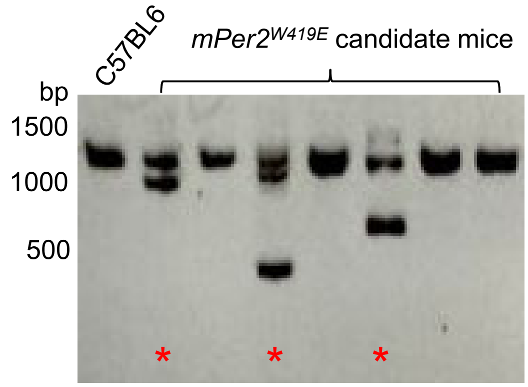


**b**

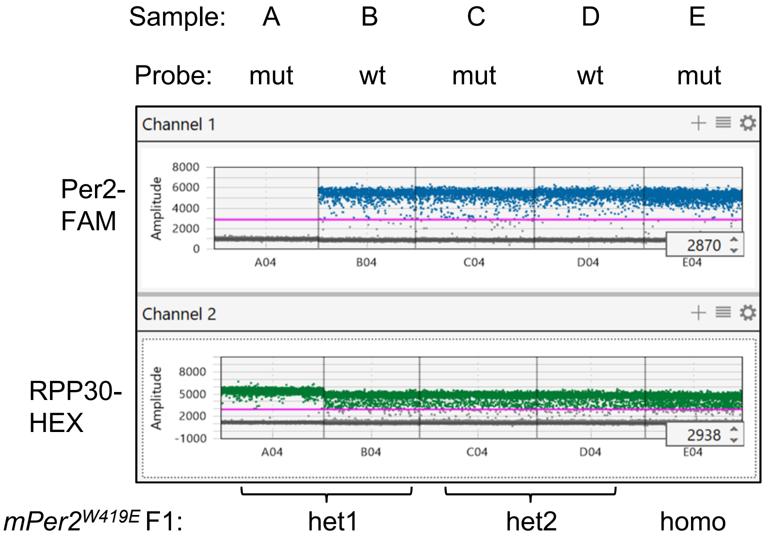


| F1 samples | Mouse ID | probe | ref | RPP30 | probe | P2W419 /RPP30 |
|---|---|---|---|---|---|---|
| A | P2 W419 #400 large del/+ | w419E probe | RPP30 | 4720 | 0 | 0.0% |
| B | P2 W419 #400 large del/+ | wt probe | RPP30 | 4832 | 2326 | 48.1% |
| C | P2 W419 #232 W419E/+ | w419E probe | RPP30 | 4732 | 2277 | 48.1% |
| D | P2 W419 #232 W419E/+ | wt probe | RPP30 | 4811 | 2345 | 48.7% |
| E | P2 W419 #402 W419E/W419E | w419E probe | RPP30 | 4945 | 4865 | 98.4% |

Fig S4

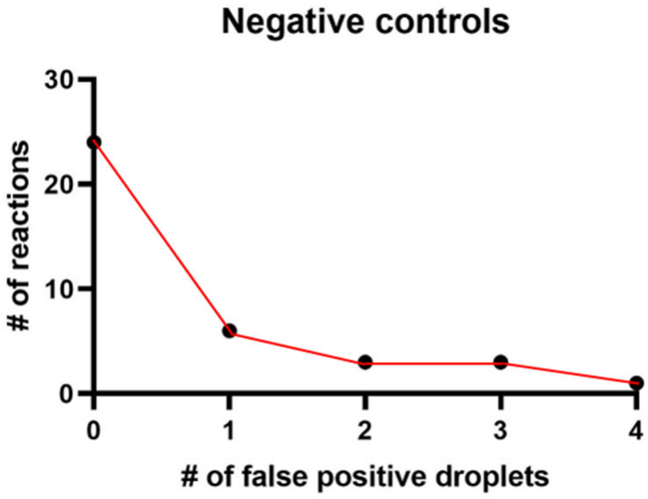
Negative controls
30
20
10
0
### of reactions
0
1
2
3
4
### of false positive droplets

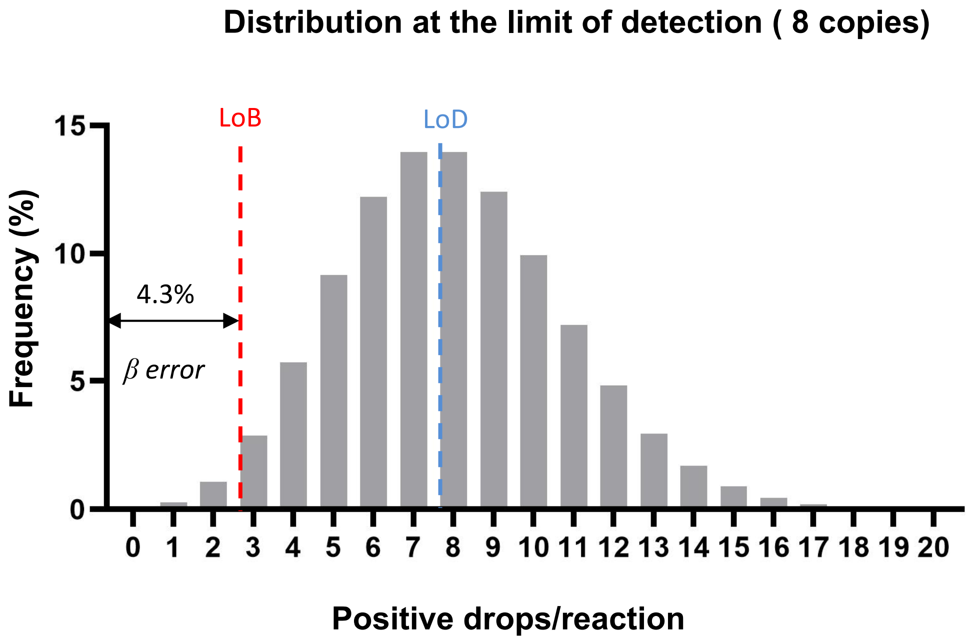
Distribution at the limit of detection ( 8 copies)
LoB
LoD
15
10
5
0
Frequency (%)
4.3%
β error
0 1 2 3 4 5 6 7 8 9 10 11 12 13 14 15 16 17 18 19 20
Positive drops/reaction
